# A Geometric Theory Integrating Human Binocular Vision with Eye movement

**DOI:** 10.1101/2020.09.03.280248

**Authors:** Jacek Turski

## Abstract

A theory of the binocular system with asymmetric eyes (AEs) is developed in the framework of bicentric perspective projections. The AE accounts for the eyeball’s global asymmetry produced by the foveal displacement from the posterior pole, the main source of the eye’s optical aberrations, and the crystalline lens’ tilt countering some of these aberrations. In this theory, the horopter curves, which specify retinal correspondence of binocular single vision, are conic sections resembling empirical horopters. This advances the classic model of empirical horopters as conic sections introduced in an ad hoc way by Ogle in 1932. In contrast to Ogle’s theory, here, anatomically supported horopteric conics vary with the AEs’ position in the visual plane of bifoveal fixations and their transformations are visualized in a computer simulation. Integrating horopteric conics with eye movements can help design algorithms for maintaining a stable perceptual world from visual information captured by a mobile robot’s camera-head. Further, this paper proposes a neurophysiologically meaningful definition for the eyes’ primary position, a concept which has remained elusive despite its theoretical importance to oculomotor research. Finally, because the horopteric conic’s shape is dependent on the AE’s parameters, this theory allows for changes in retinal correspondence which is usually considered preformed and stable.

## 1 Introduction

Our eyes receive two disparate perspective projections of a scene due to their bilateral separation. Their two-dimensional (2D) layer of photoreceptors sampling these projections is part of an unstable retinal circuitry. This happens because our eyes are constantly moving between 3-4 times per second to fixate the high-acuity fovea successively on the salient and behaviorally relevant parts of the scene [1]. Thus, there is visible motion due to eye movements even during steady fixations [2].

Therefore, the retinal images have visible motion due to both the eyes’ incessant movements and the movements of objects in the scene. Although this should lead to a compromised understanding of the scene, we instead perceive, with vivid impressions of forms in depth, stable visual scenes containing moving objects. To understand our perceived constancy of a 3D world from 2D, unstable sensory inputs, we need to understand how binocular vision is integrated with the eyes’ movements.

Whenever a retinal element is stimulated by a localized light, the stimulus is perceived in a specific direction. If the stimulus projecting to the two retinal elements, one for each eye in the binocular system, is perceived in the same direction, then they are considered to be corresponding elements. Normal correspondence occurs when the fovea of one eye corresponds to the fovea of the other eye; their single visual direction is called the principal visual direction, or the Cyclopean direction. The visual directions of all other pairs of stimulated corresponding elements are perceived in relation to this principal direction. The horopter is the set of all points in the binocular visual field stimulating retinal corresponding elements. Because the normal binocular vision is specified by two foveae being corresponding, all other corresponding retinal elements can then be determined from laboratory measurements of the empirical horopter [3, 4].

The empirical horopters were comprehensively modeled in [5, 6] and, more recently, in [7, 8]. The equations with free parameters that were introduced on an ad hoc basis in [5] for the forward gaze and extended in [6] to any horizontal gaze furnished longitudinal horopters as conic sections. Introduced in [7] and numerically studied in [8], the empirical horopters were modeled as conic sections in the binocular system with asymmetric eyes (AEs). The AE is the model eye that extends the reduced eye with its inclusion of the fovea’s displacement from the posterior pole and the cornea’s and lens’ relative tilts observed in healthy human eyes [9, 10]. This fovea’s anatomical displacement is the main source of optical aberrations and the lens’ tilts cancel out some of these aberrations by contributing to the eye’s aplanatic design [11, 12].

My studies in [8] found that the horopteric conics were numerically similar but geometrically different from the conic sections in [5, 6]; my conic sections pass through the nodal points’ anatomical location and their conic sections pass, incorrectly, through the eyes’ rotation centers. Further, in my studies, the straight-line empirical horopter, defining the abathic distance to the symmetrically fixated point, resulted from the AEs’ position in which their lens’s equatorial planes are coplanar. Then, when the AE’s parameters are set to the average values for the human eye, the resulting abathic distance of 1 meter complies with its average physiological value in humans [13]. This resulting abathic distance is also within the range of the eye muscles’ natural tonus resting position distance [14, 15].

In this paper, I extend numerical studies in [8] by developing a simple geometric theory in which the retinal correspondence of the binocular system with AEs is elaborated in the framework of bicentric perspective projections [16]. Because the eye muscles’ natural tonus resting position serves as a zero-reference level for convergence effort [17], this theory contends that the primary position of the AEs coincides with the abathic-distance bifoveal fixation. The primary position, originally intended for a single eye, is often described in binocular vision as both eyes being directed straight ahead by an erect head. This rather imprecise definition of the eyes’ primary position could be the reason for its neurophysiological significance remaining elusive despite its theoretical importance to oculomotor research [18]. Thus, this novel characterization of the eyes’ primary position integrates binocular conics with the eyes’ movements in a precise and natural way that has been unavailable until now.

The result of such an integration is that we are now able to graphically simulate the horopteric conics’ transformations from the movement of the fixation point in the visual plane, which also demonstrate the horopteric conics’ classification in terms of the eyes’ position. *GeoGebra*’s dynamic geometry software is used in this paper to demonstrate all geometric results found for the horopters and the retinal correspondence. The simulation of horopteric conics’ transformations is included in the supplementary material that is appended after the paper.

The theory’s binocular framework of bicentric retinal projections accounts for the fact that the human decodes properties of the 3D environment from neural processes fundamentally constrained by the sensory organs’ geometric relationships to the environment [19, 20]. In addition, the AE accounts for some of the eye’s aplanatic design that correlates the lens’s misalignment with corneal aberration to produce nearly diffraction-free retinal images close to the visual axis [12].

Although the distribution of retinal corresponding elements is usually considered fixed [21], the horopter’s shape and the retinal correspondence are dependent on the asymmetry parameters of the model eye and can, therefore, change when the AE’s parameters change. For example, when the crystalline lens is replaced during refractive surgery with a toric intraocular lens (IOL), it does not only correct for refractive errors and provide sharper focus, but corneal astigmatism can be also be corrected for by adjusting the lens’s orientation. The evaluation of a group of patients in [22] shows that the IOL tilt magnitude increased significantly compared to the preoperative crystalline lens’s tilt. This increase in tilt can postoperatively modify the horopter’s shape and the retinal correspondence.

## 2 Asymmetric Eye

The AE model (Figure 1), discussed in detail in [8], incorporates the most important features of the human eye’s asymmetric design. However, the AE model is slightly modified here by its use of the effective lens. The eye’s natural asymmetry is modeled by two parameters; the angle *α* which specifies the fovea’s temporalward displacement from the posterior pole, and the angle *β* which gives the effective lens’s tilt and decentration relative to the optical axis. The effective lens introduced in the AE model simplifies the description of the lens’s tilt and defines the optical axis as the eyeball’s line of axial symmetry when *α* = *β* = 0. I assume that *α − β >* 0 because it is satisfied in a typical binocular system. Because angle *α* has a low interpatient variability [23], I use the angle *α*’s average value of 5.2°. Angle *β* is assumed to vary between *−*0.4° and 4.7° and have an average value of 3.3° that conforms with the average abathic distance of about one meter.

**Figure 1:**
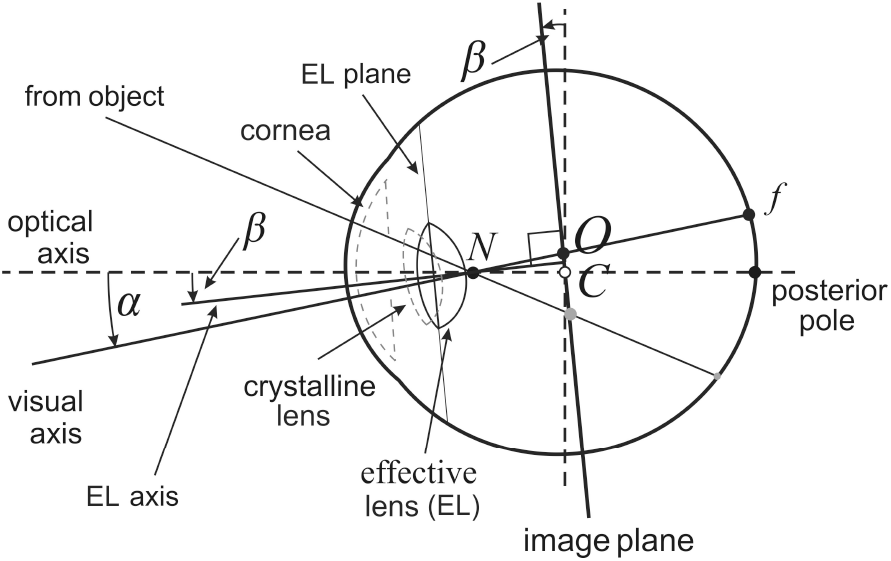
Asymmetric model eye (AE) for the right eye. The fovea, *f*, is displaced from the posterior pole by the eyeball’s global tilt of *α* degrees. The relative misalignment of the cornea and lens is represented by the *β*-degree tilt of the equatorial plane of a single effective lens. Both angles of tilt are at the nodal point *N* located on the optical axis 0.6 cm anterior to the eyeball’s rotation center *C*. The optical axis is defined by *α* = *β* = 0. The image plane is obtained by tilting the frontal plane by *β* degrees at the eyeball’s rotation center *C*. The visual axis passing through *N* and *f* intersects the image plane at its optical center *O*.

The tilt of the effective lens is represented in my geometrical model of the binocular system with AEs by the image plane passing through the eye’s rotation center that is parallel to the equatorial plane of the effective lens. The image impinged on the retina is defined by the pencil of light rays passing through the nodal point. In the AE model, these light rays may be parameterized in angular coordinates on the curved surface of the retina, or on the image plane with more convenient in image processing rectangular coordinates. The cornea and crystalline lens’s misalignment, represented by the effective lens’ tilt, is one of the eye’s sophisticated aplanatic elements designed to compensate for some of the limitations to optical quality caused by the fovea’s displacement form the eyeball’s posterior pole [12].

## 3 AEs’ Retinal Correspondence

In normal binocular vision, the foveae are corresponding elements. This means that the fixated point is perceived in one direction—the principal, or Cyclopean, direction. The horopter curve through the fixation point is the locus of spatial points that project to the retinal corresponding elements such that each point of the horopter is perceived in the same direction relative to the Cyclopean direction.

Based on the results obtained in [8], the straight line horopter shown in Figure 2, which passes through the fixation point *F*_*a*_, is established by the image planes’ coplanarity. The right AE’s visual axis passes through its respective nodal point and intersects the retina at the fovea *f*_*r*_ and the image plane at point *O*_*r*_. The other visual axis passes, similarly, through the nodal point of the left eye before intersecting the retina at *f*_*l*_ (the fovea) and the image plane at *O*_*l*_. These *O*_*r*_ and *O*_*l*_ points are the binocular correspondence centers of the image planes.

**Figure 2:**
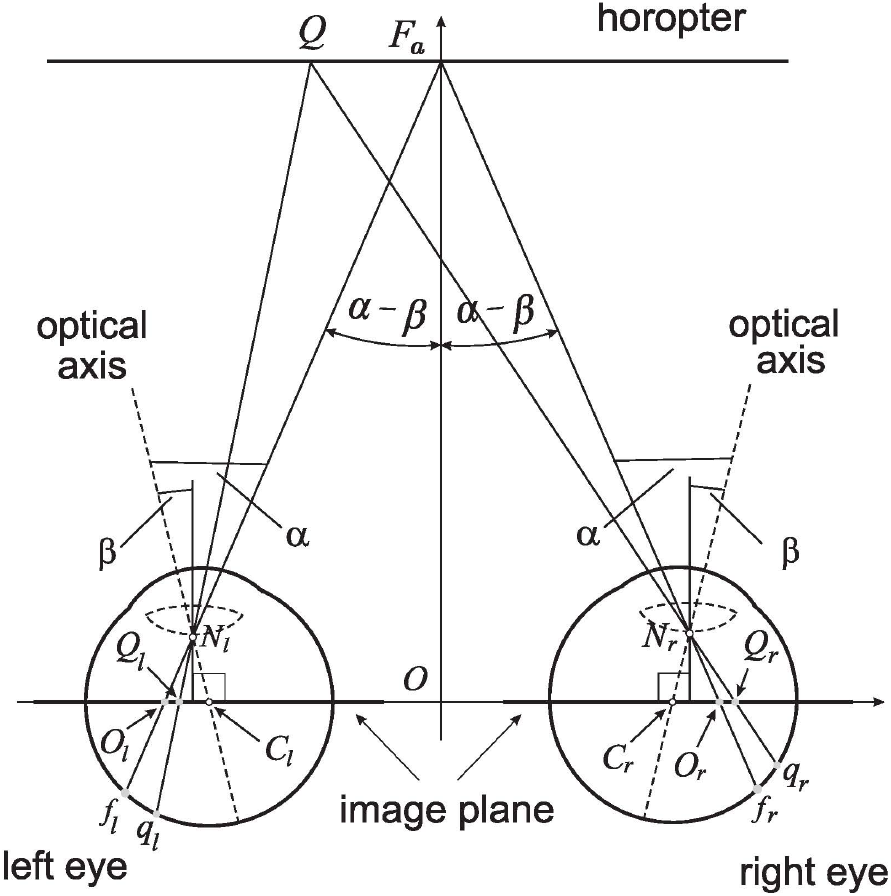
The retinal correspondence’s relation is formulated for the linear horopter at the abathic distance fixation *F*_*a*_ in the binocular system with AEs. *F*_*a*_ projects along the visual axes to the foveae *f*_*r*_ and *f*_*l*_ and the related image planes’ optical centers *O*_*r*_ and *O*_*l*_. The point *Q* projects to the retinal corresponding points *q*_*r*_ and *q*_*l*_ and their image planes’ counterparts *Q*_*r*_ and *Q*_*l*_. The asymmetric distribution of retinal corresponding points covers, under the projections through the nodal points, the symmetric distribution of related points on the image planes, which is proved in the text.

The distance *d*_*a*_ = |*OF*_*a*_| to the linear horopter at *F*_*a*_, referred to as the abathic distance, was obtained in [8]. Here, the abathic distance is given in terms of asymmetry parameters, *α* and *β*, and interocular length, 2*a* = |*C*_*r*_*C*_*l*_|, in an equivalent but simplified form,

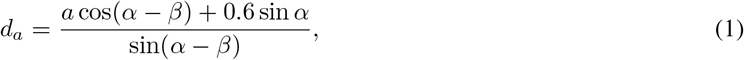

where 0.6 is the distance in centimeters from the nodal point to the eye’s rotation center.

Using the average human values 2*a* = 6.5 cm, *α* = 5.2°, and *−*0.4°*≤ β ≤* 4.7° in (1), we obtain 34 cm *≤d*_*a*_ *≤* 380 cm. However, in rare cases, the values of *β* can approach more closely the value of *α*, giving much larger value of *d*_*a*_.

A point *Q* on the linear horopter projects to retinal corresponding points: *q*_*r*_ in the right eye, *q*_*l*_ in the left eye, and *Q*_*r*_ and *Q*_*l*_ on the respective image planes, also called the image planes’ corresponding points. The corresponding points *q*_*r*_ and *q*_*l*_ are located at different distances from their respective foveae such that the asymmetric distribution of corresponding retinal points with respect to the foveae is the result of the eyes’ asymmetry and the head’s bilateral symmetry. However, from similar triangles, *QF*_*a*_*N*_*r*_ and *Q*_*r*_*O*_*r*_*N*_*r*_ for the right eye, and similar triangles, *QF*_*a*_*N*_*l*_ and *Q*_*l*_*O*_*l*_*N*_*l*_ for the left eye, we conclude that |*Q*_*r*_*O*_*r*_| = |*Q*_*l*_*O*_*l*_|. This line horopter and the head’s bilateral symmetry [24, 25] is used here to define the binocular correspondence as follows.

**Retinal Correspondence**. *Referring to Figure 2, let O*_*r*_ *and O*_*l*_ *be projection points of the abathic-distance fixation point F*_*a*_ *into the right and left image planes of AEs, respectively. Then, any two points Q*_*r*_ *and Q*_*l*_ *of the equal distance from, and on the same side of O*_*r*_ *and O*_*l*_ *project, through the nodal points N*_*r*_ *and N*_*l*_, *to the retinal corresponding points q*_*r*_ *and q*_*l*_ *of unequal distance from the foveae f*_*r*_ *and f*_*l*_, *respectively*.

I demonstrate later in Remark 1 of Section 5 that the binocular correspondence defined by the line horopter is indeed a well-defined concept, i.e., it does not depend on the eyes’ bifoveal fixation. This definition of retinal correspondence has previously only been assumed in [8]. However, the distribution is not homogeneous because the retinal photoreceptors are not uniformly distributed on the retinae; instead, they are concentrated in the foveal areas. The binocular correspondence is only specified up to the ratio of the symmetric distribution of corresponding points on the image planes to the asymmetric distribution of corresponding points on the retinae. A precisely specified binocular correspondence’s relationship can only be obtained for each subject in laboratory measurements.

## 4 Binocular System with AEs

This section introduces basic definitions in the geometry of the binocular system with AEs displayed in Figure 3, but a detailed elaboration of binocular geometry is developed throughout the following sections. In particular, the horopteric curves are geometrically constructed in the next section.

**Figure 3:**
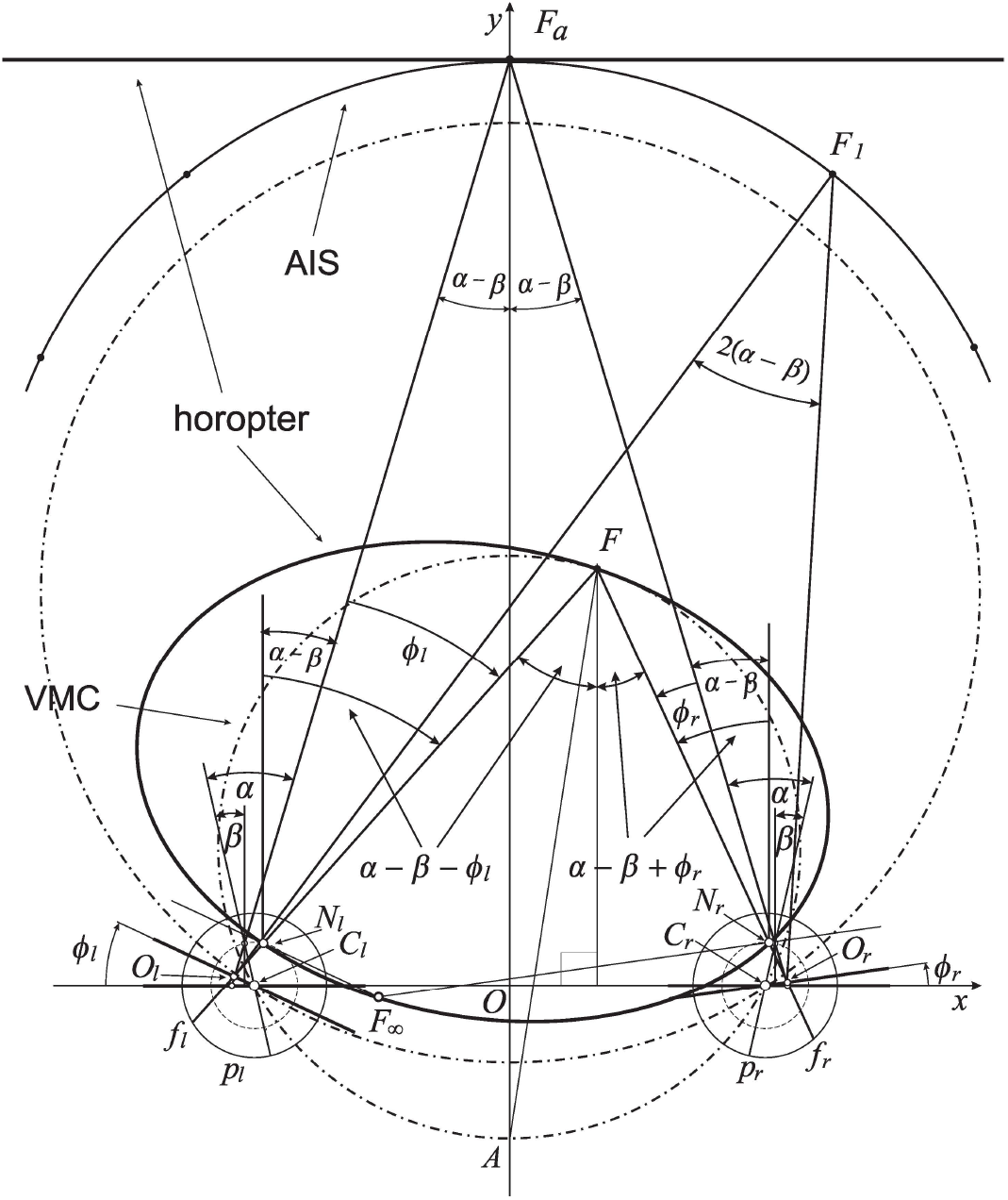
The eyes’ asymmetry angles *α* and *β* are shown only for fixation *F*_*a*_. The rotating angles, *ϕ*_*r*_ and *ϕ*_*l*_, change the eyes’ gaze from the resting vergence position, fixation *F*_*a*_ at the abathic distance, to the position in which eyes are fixating on *F*. This results in the subtense changing from *σ*_*a*_ = 2(*α − β*) at *F*_*a*_ to the subtense *σ*_*F*_ = 2(*α − β*) + *η* at *F*, where *η* is the vergence (2). The horopteric ellipse, shown here for the fixation *F*, is constructed in Section 5 using the nodal points, *N*_*r*_ and *N*_*l*_, and the intersection point, *F*_*∞*_, of the lines through the nodal points and parallel to the respective image planes. The condition *ϕ*_*r*_ *− ϕ*_*l*_ = 0 furnishes a curve with a constant subtense *σ*_*a*_. This is the abathic iso-subtense curve (AIS) that passes through *F*_*a*_. Later, similarly to the case of the symmetric (reduced) model eye, the Cyclopean direction of the fixation point *F* in the binocular system with AEs will be specified relative to the point *A* on the VMC passing through *F*.

For the average human eye’s asymmetry parameters *α* = 5.2° and *β* = 3.3°, and the average ocular separation 2*a* = 6.5 cm, the abathic distance (1) is 99.61 cm, a value consistent with the average value recorded in humans [13]. This distance is similar to the eye muscle’s natural tonus resting position distance [15], which serves as a zero-reference level for the eyes’ convergence effort [17]. Therefore, I refer to the position of the eyes fixating at the abathic-distance as the resting vergence position, in order to distinguish it from the primary eyes’ position, often described as both eyes directed straight ahead in an erect head.

For the average human eye’s asymmetry parameters *α* = 5.2° and *β* = 3.3°, and the average ocular separation 2*a* = 6.5 cm, the abathic distance (1) is 99.61 cm, a value consistent with the average value recorded in humans [13]. This distance is similar to the eye muscle’s natural tonus resting position distance [15], which serves as a zero-reference level for the eyes’ convergence effort [17]. Therefore, I refer to the position of the eyes fixating at the abathic-distance as the resting vergence position, in order to distinguish it from the primary eyes’ position, often described as both eyes directed straight ahead in an erect head.

I note that in the binocular system with symmetric eyes, i.e., with model eyes satisfying *α* = *β* = 0, *ϕ*_*r*_ and *ϕ*_*l*_ are angles describing the eyes’ rotations from their primary position. In this case, the angle subtended at the resulting fixation point is given by the vergence angle

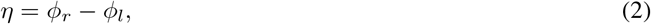

cf. fixation *F*_0_ in Panel C of Figure 4.

**Figure 4:**
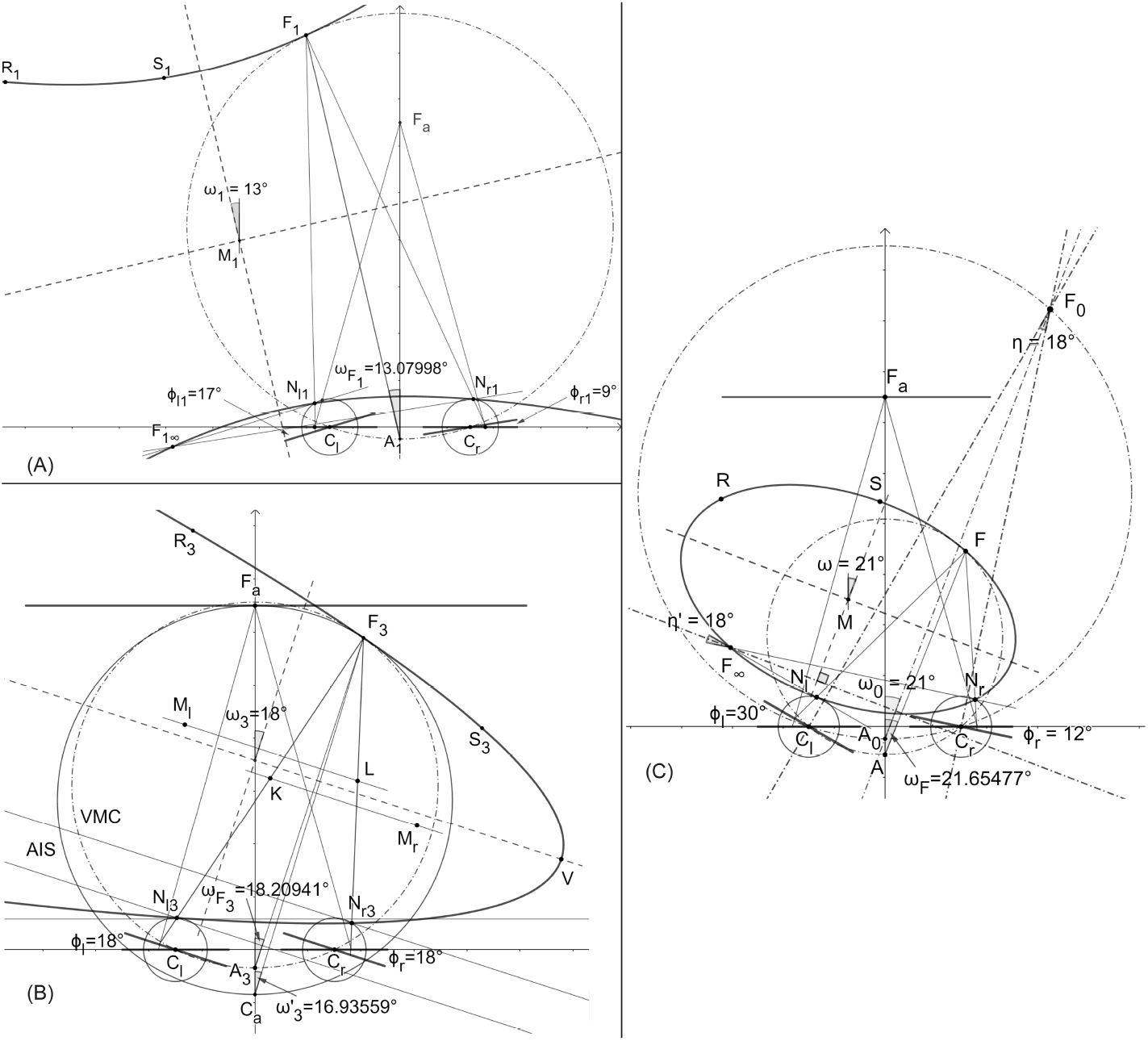
The horopteric conic sections constructed in the demonstration of Binocular Conics Construction. Panel A. The hyperbola. Panel B. The parabola. Panel C. The proof that the conic section orientation is given by version angle, carried out only for the ellipse.

To get the subtense of the fixation points *F*_*a*_ and *F*, I use the equality of alternate angles: two angles, not adjoined, formed on opposite sides of a line where the line intersects two other parallel lines. In Figure 3, *α − β − ϕ*_*l*_ at both vertices, *N*_*l*_ and *F*, are alternate angles for the left eye, and *α − β* + *ϕ*_*r*_ at both vertices, *N*_*r*_ and *F*, are alternate angles for the right eye. Note that the angle *ϕ*_*l*_ is subtracted from *α − β* because its value is negative. It is easy then to verify, by taking the sums of respective alternate angles, that the angle subtended by visual lines at *F*_*a*_ is 2(*α − β*) and the angle at *F* is 2(*α − β*) + *η*. Thus, since *α ≠ β*, the angle at any fixation obtained by the change of gaze from *F*_*a*_ never takes on the vergence angle *η* in (2). Therefore, in this work, the angles subtended by the visual lines at the spatial points are called binocular subtense, or just subtense.

Eye positions reached from the resting vergence position by equal eye rotations, *ϕ*_*r*_ = *ϕ*_*l*_, have fixation points that lie on, what I call, the abathic iso-subtense curve (AIS). For each different symmetric fixation of subtense 2(*α − β*) + *η*, we get different iso-subtense curve. This curves differ from the iso-vergence curves, or Vieth-Müller circles (VMCs), because, in contrast to the iso-subtense curves, the VMC passes through the eyes’ rotation centers. The AIS curve, the iso-subtense curve which passes through *F*_*a*_ at the abathic distance, is graphed numerically in Figure 3 for fixations in the azimuthal range *±* 45°, the neurally determined range of typical gaze eccentricities [26]. For anthropomorphic binocular system parameters, the AIS will be closely approximated in Section 6 by the VMC.

Further, for symmetric eyes (*α* = *β* = 0), the version angle,

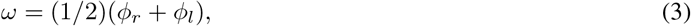

is the azimuthal angle of the ray that starts from the point on the VMC’s that is midway between the eyes’ centers and passes through the fixation point (cf. the fixation point *F*_0_ in Panel C of Figure 4) given by the rotation angles *ϕ*_*r*_ and *ϕ*_*l*_ from the eyes’ primary position. The VMC’s midpoint and the azimuthal angle (3) specify the Cyclopean eye’s position [27] and, hence, the principal visual direction. Section 6 will discuss how the fixation point *F* ‘s Cyclopean eye position can be defined in the binocular system with AEs.

## 5 The Geometric Construction of Binocular Conics

In the binocular system with AEs, parameters *α, β, a* and the eyes’ rotation angles *ϕ*_*r*_ and *ϕ*_*l*_ specify coordinates of the four points in the horizontal visual plane which lie on, or are associated with, the corresponding horopteric curve. These points are the nodal points *N*_*r*_ and *N*_*l*_, the fixation point *F*, and the point denoted by *F*_*∞*_. The point *F*_*∞*_ (cf. Figure 3) is the intersection of the two lines, each passing through the nodal point of one eye and parallel to both the AE’s image plane and effective lens’s equatorial plane. Thus, in the projective geometry framework [28], *F*_*∞*_ projects to the points at infinity, one for each of the AE’s image planes. The fixation point *F* projects along the visual axes to the foveae, which are corresponding retinal elements in normal binocular vision. In contrast, the lines projected from *F*_*∞*_ to the pair of points at infinity do not intersect the retinae.

Thus, the points at infinity are not corresponding retinal elements though they are called geometrical corresponding points here because of the significant role they play in the horopteric curves’ geometric constructions. These geometrical constructions for the binocular system with AEs are given below in this section. These constructions are motivated by the results obtained in [27] for horopteric circles in binocular system with symmetric (reduced) eyes. They are reframed here in Proposition 1 to include *F*_*∞*_ into the formulation, which is otherwise not needed because it is a theorem of Euclidean geometry.

### Proposition 1.

*Let the nodal point be located on the optical axis at any point at or between the spherical eyeball’s rotation center and its pupil. Then, for the binocular eyes’ position with fixation point F in the horizontal visual plane, the lines which pass through the nodal points and are perpendicular to the visual axes intersect at the point F*_*∞*_ *on the circular horopter. It then follows that line segment FF*_*∞*_ *must pass through the horopteric circle’s center*.

The proof of Proposition 1 is given in Supplement 1. It shows that, in the binocular system with symmetric model eyes, *F* and *F*_*∞*_ are diagonally opposite points on the horopteric circle. The anatomically correct location of the nodal point is 0.6 cm anterior to the eye’s rotation center, though the proof is for any nodal point location between the eye’s rotation center and pupil.

The construction of horopteric curves in binocular system with AEs incorporates the horopteric circles’ point symmetry of Proposition 1. The rationale for this extension is the continuity requirement of the horopteric curves’ transformations as the AE’s parameters *α* and *β* both approach zero. Moreover, referring to my previous research, the extension also accurately reflects *F* and *F*_*∞*_ ‘s relation in projective geometry, the geometric framework which is essential to the constructions of horopteric curves for the binocular system with AEs.

To explicate this further, I note that the mapping between points of the spherical retina and points of the image plane can be modeled by stereographic projection through the nodal point for both symmetric and asymmetric model eyes [7]. This mapping is not defined at the nodal point. Stereographic projection is extended to one-to-one and onto by appending the image of the nodal point under the mapping, called the point at infinity, to the image plane. The image plane with the point at infinity is the celebrated object in geometry and mathematical analysis known as the Riemann sphere, [29]. Stereographic projection is conformal, that is, it preserves the angle of two intersecting curves. Further, it maps circles in the spherical retina that do not contain the nodal point to circles in the image plane. Therefore, this conformal geometry preserves receptive fields and retinal illuminance, providing constructive properties for human vision [30].

Now, for each of the binocular system’s AEs, the fixation point *F* in the horizontal visual field defines the origin in the image plane and *F* _*∞*_is projected to the point at infinity. The origin and the point at infinity are images of the fovea and the nodal point under stereographic projection which identifies the spherical retina with the image plane and, therefore, they are opposite points in the Riemann sphere. I assume that *F* and *F*_*∞*_ are opposite points on the horopter of the binocular system with AEs. This assumption, which is confirmed in this paper by geometric constructions supported with dynamic geometry software, provides us with a particularly simple theory of empirical horopters that is both biologically supported and geometrically precise, advancing the classic model of empirical horopters introduced by Ogle in [5]. Surprisingly, both stereographic projection and the horopter were first introduced by Aguilonius in his Six Books of Optics published in 1613.

The demonstration of the main results of horopteric conics, referred to as binocular conics, is constructive and, thus, making possible to design algorithms for modeling stable binocular vision in mobile robots.

**Binocular Conics Construction.** *For the binocular system with AEs’ orientations such that point F_∞_ is in the visual field, the horopteric curve’s center is designated the midpoint M of line segment FF_∞_. This means that for each point on the curve, there is another point on this curve diagonally opposite to it. Then, this curve, either an ellipse or a hyperbola, is fully specified by F, F_∞_ and the nodal points N_r_ and N_l_. Further, when F_∞_ is at infinity, that is, when the image planes are parallel but not coplanar, the horopteric curve is a parabola. Each conic section’s orientation is exactly given by the version angle (3)*.

Demonstration: The horopteric curves in the binocular system with AEs are binocular conics under the assumed point symmetry of the horopteric circles in Proposition 1, they are geometrically constructed and graphically visualized in *GeoGebra* (Figure 4). Because the construction involves the same steps for hyperbolas and ellipses, I construct only a hyperbola. For a given position in which the eyes’ image planes are non-parallel, the eyes’ nodal points *N*_*r*1_ and *N*_*l*1_, fixation point *F*_1_, and point *F*_1*∞*_, are all shown in Panel (A) of Figure 4. Then, two additional points on the conics are constructed in Panel A by taking reflections of the nodal points about midpoint *M*_1_ of line segment *F*_1_*F*_1*∞*_. These additional points, shown as *R*_1_ and *S*_1_, determine the conics. Shown in this panel, the conic section graphed in *GeoGebra* by taking any five of these six constructed points, is the same hyperbola. Symmetric fixation at the abathic distance has coplanar image planes. For the abathic distance fixation, the fronto-parallel linear horopter was constructed in Section 3. For any fixation obtained from *F*_*a*_ by the same rotation angle of both eyes, the resulting image planes are parallel but not coplanar and *F*_*∞*_ is at infinity. In the projective geometry framework, *F*_*∞*_ is represented by a family of lines parallel to the eyes’ image planes and the conics are parabolas. One of these parabolas is constructed for fixation *F*_3_ in Panel B as follows. First, midpoint *L* of the line segment connecting *N*_*r*3_ and *F*_3_ is obtained and the line in the visual plane through *L* that is parallel to the image planed. This line intersects the line that passes through *N*_*l*3_ and is parallel to segment *N*_*r*3_*F*_3_ at the point *M*_*l*_. Then, point *R*_3_ on the parabola we want to construct is obtained by reflecting *N*_*l*3_ about the point *M*_*l*_. The same steps are repeated starting with line segment *N*_*l*3_*F*_3_ to obtain points *K* and *M*_*r*_. These points then give the second point *S*_3_ on the parabola by taking a reflection of *N*_*r*3_ about *M*_*r*_. All steps in these constructions, for example, taking a reflection about a point, are done via dynamic geometry environments of the *GeoGebra* system. The proof of the conic sections’ orientations, specified by the version angle (3), is given for the horopteric ellipse in Panel C of Figure 4. The rays passing through *F*_*∞*_intersect at 18°. This is the vergence at fixation point *F*_0_ that is obtained for the symmetric eyes’ azimuthal rotation angles *ϕ*_*r*2_ = 12° and *ϕ*_*l*2_ = 30°. The two bisecting lines at *F*_*∞*_ and *F*_0_ intersect at the right angle, proving that the orientation of the ellipse is given by the angle *ω* = 21° equal to the version *ω*_0_ = 21°. This proof also holds for hyperbolas and parabolas. Because the values of the eye’s asymmetry parameters are chosen arbitrarily, this proof applies to any parameters chosen for the eyes. Thus, the link between the horopter’s geometry and eye movements is established. This ends DEMONSTRATION.

The geometrical construction of binocular conics in *GeoGebra* allows the graphical simulation the horopteric conics’ transformations from the movement of the fixation point in the visual plane. The computer simulation of horopteric conics’ transformations is included in Supplement 2.

I recall that the binocular correspondence was defined in Section 3 using the straight-line horopter for the symmetrically fixated point at the abathic distance. This is a well-defined concept only if the corresponding retinal points are independent of the binocular conics transformations when the fixation point moves in the visual plane.

Each point on one of the binocular conics projects along the eyes’ visual lines to the retinae of the AEs and defines one pair of corresponding points. However, only two pairs of points are used in the binocular conics’ construction: the two foveae and the two points at infinity. All other corresponding retinal elements are established by the bicentric retinal projections of the horopter’s points. In a computer simulation in the *GeoGebra*, available in Supplement 2 A and B, retinal corresponding points, *q*_*r*_ and *q*_*l*_, and the image plane’s corresponding points, *Q*_*r*_ and *Q*_*l*_, are both determined by point *Q* (cf. Figure 2) on the abathic distance line horopter and so remain corresponding when the eyes’ position changes in the visual plane of fixations. Because this must be true for all pairs of retinal corresponding points, I conclude that

### Remark 1.

*The Binocular correspondence’s relationship introduced in* **Retinal Correspondence** *in Section 3 is well-defined*.

How are these intrinsic properties of the theory related to human binocular vision? The human brain functions in physical space and receives information carried by light that is centrally projected onto the eyes’ retinae and transduced by photoreceptors into electrochemical signals. After initial processing by the retinal circuitry, this visual information is mainly sent to the primary visual cortex where it produces specific retino-cortical mappings and forms input to other cortical areas [31]. This immensely complex processing decodes the environment from retinal stimulation and creates a neural representation of space [32], our subjective visual space.

The newest computational modeling in neuroscience that incorporates bicentric perspective mapping of the 3D environment onto the retinae demonstrates that this mapping is fundamental to the tuning of retino-cortical neuronal processes and these process’s corresponding aspects of perception [20]. Although the tuning was specifically examined for 3D motion in the primate cortical area MT, the process of decoding the world from retinal stimulation in visuomotor cortical areas must be strongly affected by the geometry that links the environment to the sensory epithelium, regardless of whether nonhuman or human primates locomote or scan the environment while standing still. This geometric relationship constraining visual perception in my theory is the bicentric projective mapping between 3D space and the AEs’ image planes which are determining the horopter’s shape. The horopter’s shape in turn establishes a well-defined retinal correspondence. This theoretical relationship mirrors the one in human binocular vision in which retinal correspondence of normal binocular vision is specified by the two, corresponding, foveae such that all other corresponding retinal elements are then determined from laboratory measurements of the empirical horopter. However, the question of whether the corresponding retinal elements are fixed or not has remained undecided [21, 33].

In the theory presented here, binocular correspondence depends on the eye’s asymmetry parameters. Therefore, the retinal correspondence can change when the asymmetry parameters change. Such changes can occur during refractive surgery. For example, to correct for refractive errors and achieve sharper vision, which is common for people with presbyopia, the crystalline lens are surgically replaced with an artificial intraocular lens (IOL). Toric IOLs can also correct astigmatism caused by the shape of the cornea by adjusting the lens’s orientation because they have different powers in different meridians. When a group of 333 patients were evaluated in [22] for preoperative crystalline lens and postoperative IOL tilt, their IOL’s tilt magnitude was found to have increased significantly by 1.2° *±* 1.1° compared to the preoperative crystalline lens tilt. I conclude from these results that postoperative change in the lens’ tilt can be large enough to change the patient’s empirical horopter’s shape and the horopter’s retinal correspondence. In the binocular system with asymmetric eyes, this change in the lens’s tilt is modeled by the angle *β*’s corresponding change.

## 6 Anthropomorphic Binocular Conics

Figure 5 depicts the binocular conics given by the numerical method from [8] (dashed lines) and the geometric method developed in Section 5 (solid lines) and drawn by *GeoGebra*’s software for the average parameters observed in humans: *α* = 5.2°, *β* = 3.3°, and ocular distance 2*a* = 6.5 cm. From the figure, we see that the hyperbolas for fixation *F*_1_ and the ellipses for fixation *F*_2_ obtained by both methods overlay each other nearly perfectly. However, the horopteric parabola for fixation *F*_3_ on AIS differs from the tangent line to the VMC at *F*_3_. Figure 5 shows that the difference between AIS (solid line through *F*_*a*_) and VMC (dot-dashed line through *F*_3_) should be insignificant to the perceptually important 90° of the central visual field.

**Figure 5:**
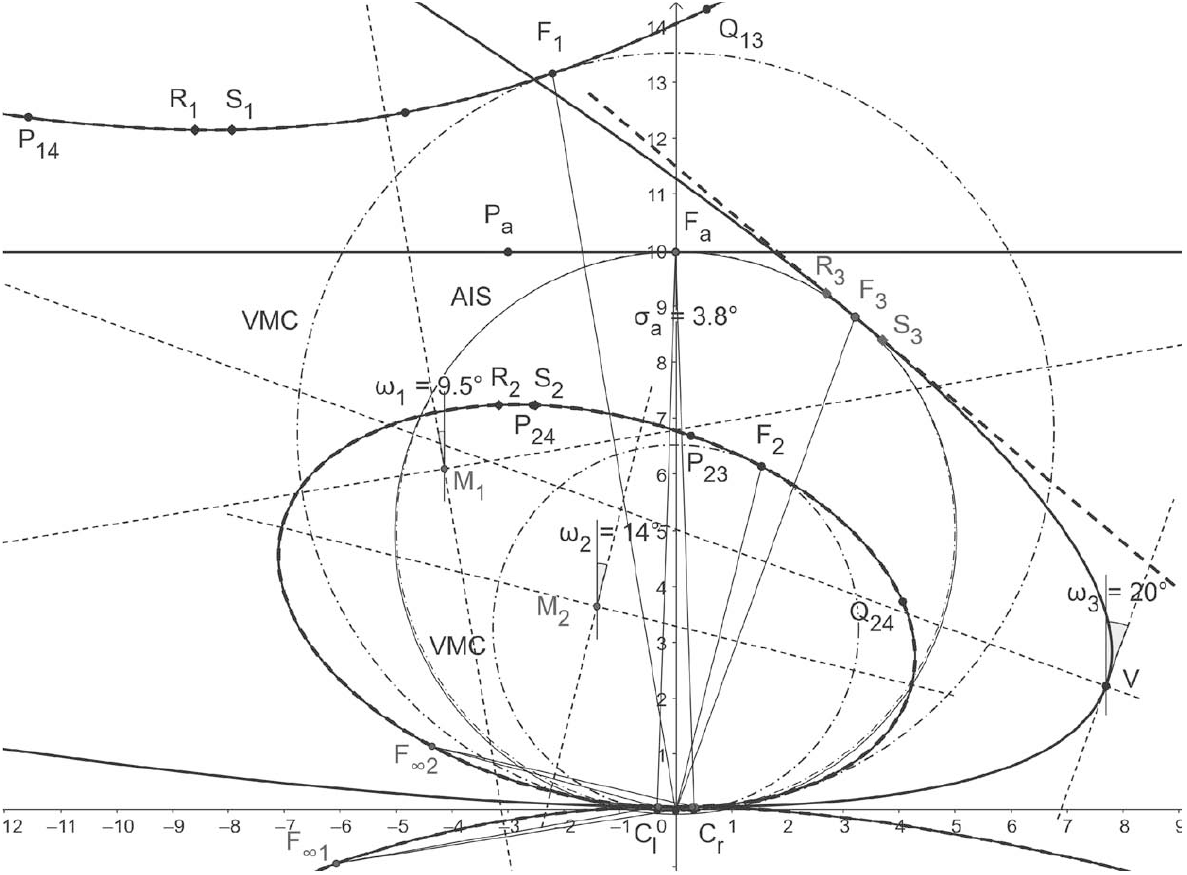
Horopteric conics for anthropomorphic parameters. The conics shown in solid lines: the line horopter for *F*_*a*_ at the abathic distance of 99.61 cm, the hyperbola for fixation *F*_1_, the ellipse for fixation *F*_2_, and the parabola for fixation *F*_3_ on the iso-subtense curve AIS (solid line) through *F*_*a*_ are constructed in Section 5. Each fixation point is on the corresponding VMC (dot-dash line). The conics obtained for the same fixation points by the method from [8] are shown in dashed lines. As we can see, the hyperbola and ellipse for both methods overlay each other nearly perfectly. The difference between the parabola for *F*_3_ and the tangent line to the VMC at *F*_3_ is explained in the next section. The conics’ orientations are given by the version angles in **Binocular Conics Construction** of Section 5.

To find the difference between the AIS and VMC, I first note that the AIS can be well approximated with a circle. In fact, using *GeoGebra*, I find that the AIS’s approximation in the visual field’s range of *±*45° (cf. Section 4) to 2 decimal places is the circle *x*^2^ + (*y −* 49.46)^2^ = (50.13)^2^. To find the equation of the VMC passing through *F*_3_, I recall the exact geometric description of the VMC given in [27]: the center (0, *k*) = (0, *a/*(2 tan *η*) and the radius *R* = *a/*(2 sin *η*). Then, upon substituting *a* = 3.25 cm and using the computed in *GeoGebra* value of *η* = 3.73° at *F*_3_, the VMC’s equation is *x*^2^ + (*y −* 49.80)^2^ = (49.91)^2^. This verifies that the difference between the two circles is negligible.

In Figure 5, all angles are obtained in *GeoGebra* by the geometric method of this paper and displayed to an accuracy of 5 decimal points. The fixation point *F*_1_ is obtained by rotations *ϕ*_*r*_ = 9° and *ϕ*_*l*_ = 10° from the resting vergence position and gives rise to the hyperbola constructed by Proposition 5.2’s method. Version *ω*_1_ = 9.5° gives the hyperbola’s orientation while *F*_1_’s orientation angle is 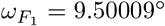. The fixation point *F*_2_ results from rotations by *ϕ*_*r*_ = *−*13° and *ϕ*_*l*_ = *−*15° and gives rise to the ellipse, again with an orientation specified by version *ω*_2_ = *−*14° and the direction of *F*_2_, which was given by azimuthal angles 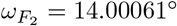;. Rotations *ϕ*_*r*_ = *ϕ*_*l*_ = *−*20° from the resting vergence position gives the fixation point *F*_3_ on AIS and version *ω*_3_ = *−*20° specifies the resulting parabola’s symmetry axis’s direction. The point on the corresponding VMC midway between the eyes provides the direction, 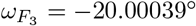, of *F*_3_. We see that the conics’ orientations and the fixation points’ directions differ by less than 4 seconds of an arc. This insignificant difference allows me to place the Cyclopean eye at the same point on the corresponding VMC it was placed at in the binocular system with symmetric eyes [27]: midway between the eyes’ centers.

We conclude from Figure 5 and the results in [8] that, for the anthropomorphic binocular system, the binocular conics in my theory are numerically close to the conic sections obtained in [5, 6]. I can therefore express the conic sections’ parameter *H* used in those studies in terms of the AE’s parameters *α* and *β*. At the abathic distance, *H* = 2*a/d*_*a*_, 2*a* = 6.5 cm is the interocular separation and *d*_*a*_ is the abathic distance to the fixation point given in (1). Thus, *H* can be expressed in terms of the eye’s asymmetry parameters as follows:

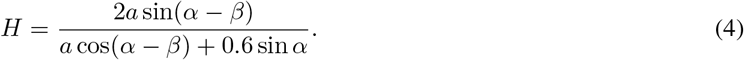

Because the subtense *σ*_*a*_ = 2(*α − β*) at the abathic distance fixation is small (0.066 radians for the anthropomorphic parameters *α* = 5.2°, *β* = 3.3°), we obtain the approximation *H ≈* 0.065 which differs from *σ*_*a*_ by approximately 0.001 rad. Moreover, for *α* = 5.2° and the range of *β*’s values, *−* 0.4° *< β <* 4.7°, cf. Section 2, I obtain the range of *H*’s values in (4) as follows: 0.01 *< H <* 0.19. This result for *H*’s values is consistent with Ogle’s original estimation of 0 *< H <* 0.2 for human subjects [5, 3]. In [3], Ogle presented the values of *H* calculated from the data of Helmholtz, Lau and Libermann among many other researchers obtained in Nonius observations, which are in the range of his original values of *H* reported in [5]. See also the relevant discussion in [4]. It is also consistent with the values estimated more recently in [34, 35]. However, these recent studies were more general by considering the Hering-Hillebrand deviation parameter *H* and the Helmholtz shear, or the vertical horopter’s backward inclination, that is not included in my study.

## 7 Binocular Conics in Visual Plane

A theory of horopteric circles in the binocular system with symmetric (reduced) eyes may be based on Euclidean geometry alone. But for a theory of horopteric conics in the binocular system with AEs, a framework of projective geometry is necessary. In projective geometry terms [28], the general conic equation given by the inhomogeneous quadratic polynomial *c*(*x, y*),

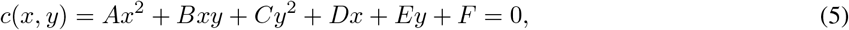

is also expressed by the homogeneous quadratic form *C*(*X, Y, Z*) = *Z*^2^*c*(*X/Z, Y/Z*).

Although no more than five points on a conic are needed to find its equation, this straightforward task appears computationally unfeasible for binocular conics because the expressions for the points specifying a generic binocular conic are too complicated. To circumvent this limitation, I classify binocular conics in terms of (5)’s discriminants and analyze the classes of conics in the ‘general position’ when the point of bifoveal fixation moves in the horizontal visual plane. The notion ‘general position’ will be explained below in this section.

The conic (5) is degenerate if and only if its discriminant, i.e., the determinant Γ of the symmetric matrix of its homogeneous quadratic polynomial, vanishes. Here,

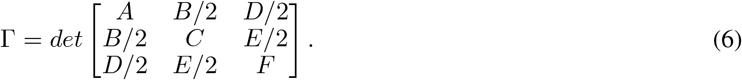

Then, for either degenerate or nondegenerate conics, its type is determined by the sign of the quadratic part of (6)’s discriminant,

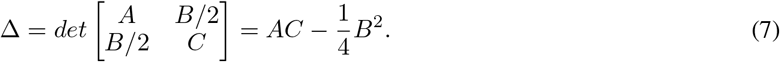

The cases restricted to the real degenerate conics Γ = 0 are: two intersecting lines Δ *<* 0, two parallel lines Δ = 0, and one point Δ *>* 0. The nondegenerate conics Γ ≠ 0 are classified as follows: the conic is a hyperbola if Δ *<* 0, an ellipse if Δ *>* 0, and a parabola if Δ = 0, see [36].

The three conics: the hyperbola for the fixation *F*_1_, the ellipse for the fixation *F*_2_, and the parabola for the fixation *F*_3_, shown in Figure 5, have the following equations and discriminants (7) obtained in the numerical simulations with *GeoGebra* for the calculated points in the constructions carried out for demonstration of Binocular Conics Construction in Section 5.

1. Hyperbola’s branch containing *F*_1_: 2.73*x*^2^ + 3.77*xy −* 8.22*y*^2^ *−* 0.40*x* + 115.86*y −* 7.11 = 0; Δ = *−*26
2. Ellipse containing *F*_2_: 1.121*x*^2^ + 0.90*xy* + 2.81*y*^2^ *−* 0.12*x −* 19.24*y* + 0.99 = 0; Δ = 2.9
3. Parabola containing *F*_3_: 0.21*x*^2^ + 1.14*xy* + 1.56*y*^2^ *−* 0.14*x −* 17.72*y* + 0.97 = 0; Δ = *−*6 *×* 10^*−*7^ *≈* 0

The discriminant value for the parabola, which should be 0, is only approximated by *−*0.0000006. This explains why the parabola was approximated in [8] by a straight line whereas, here, it is given by the parabola, cf. Figure 5. In this theory, the parabola is built into the model of horopteric curves by way of construction. However, in the numerical simulation in [8], the conics’ sensitivity near Δ = 0 allows us to see either an ellipse or hyperbola with the shape that resembles the tangent line near the fixation point. What could explain this?

Intuitively, the set of conics satisfying condition Γ = 0 is negligible when compared to the set of conics satisfying Γ ≠ 0 because the number of conics enumerated by *{*Γ = *x, x ∈ R \{*0*}}* is huge compared to conics enumerated by Γ = 0. Similarly, when Γ ≠ 0, the set of conics satisfying Δ = 0 is negligible when compared to the set of conics satisfying Δ ≠ 0. In mathematics, see [37] for example, the ‘general position’ is a notion of genericity for geometric objects satisfying some special conditions that distinguishes them from all other geometric objects in a given collection. Thus, in the whole collection, the subcollection of objects in their general position is ‘massive’, and the complementary set ‘meager’, with its objects ‘negligible’. Thus, only ellipses and hyperbolas are conics in the general position.

Now, after these preliminary remarks, I can analyze the binocular conics in the visual plane of bifoveal fixations. To this end, I note that the fixation points inside the AIS curve produce the eyes’ positions such that *η* = *ϕ*_*r*_ *− ϕ*_*l*_ *>* 0, while the eyes’ positions with fixation points outside the AIS satisfy *η* = *ϕ*_*r*_ *− ϕ*_*l*_ *<* 0. This simple property of the eyes’ positions and the computer simulation of binocular conics lead to the following proposition about horopteric geometry in the visual plane:

**Binocular Conics Transformation.** *If α > β, then the AIS of a constant subtense of* 2(*α − β*) *divides the visual plane into three distinct regions: (A) The fixation point F on the AIS determines the horopter as a parabola if F* = *F*_*a*_, *and a straight line if F* ≠ *F*_*a*_. *(B) The fixation point outside of the AIS specifies the branch of a horopteric hyperbola through this point, possibly degenerating into two intersecting lines at some of the fixation points. (C) The fixation point in the binocular field inside of the AIS specifies a horopteric ellipse. On the other hand, in the monocular field inside the region enclosed by AIS, an ellipse can change into a hyperbola such that the sequence of transformed conics passes through the degenerate case of two parallel lines*.

This classification of binocular conics transformations is demonstrated in a *GeoGebra* simulation for human-like binocular system parameters when the fixation point moves in the visual plane (Supplement 2 A).

This simulation provides the binocular conics ‘noisy’ classifications given in terms of eye position, information that is available to the visual system. When the fixation point is moved in the visual plane, only the initial linear horopter and the evolution of subsequent conics in the general position can be observed, hence the term ‘noisy’. For example, the degenerate conics cases mentioned above in **Binocular Conics Transformation**: the two intersecting lines and two parallel lines, can only be inferred from observing neighboring conics in their general positions. The three typical cases of observed binocular conics: the straight line as the initial horopter at the abathic distance fixation, and the hyperbola and ellipse obtained in the simulation, are shown, respectively, in Figure 6’s panels (A), (B) and (C). Atypical cases of the simulation are also shown in Figure 6 in panels (D), (E) and (F). In these panels, the conic morphs through two parallel lines from ellipses to hyperbolas. Further, in panel (E), the conic shown is indeed a hyperbola, although its branches may appear in the figure to be parallel lines. Running several dozens of simulation sessions attests that this will only happen well outside the binocular region. In these three last panels, points where the visual axes intersect are near the eyeball in a space where fixations are prevented by human anatomy.

**Figure 6:**
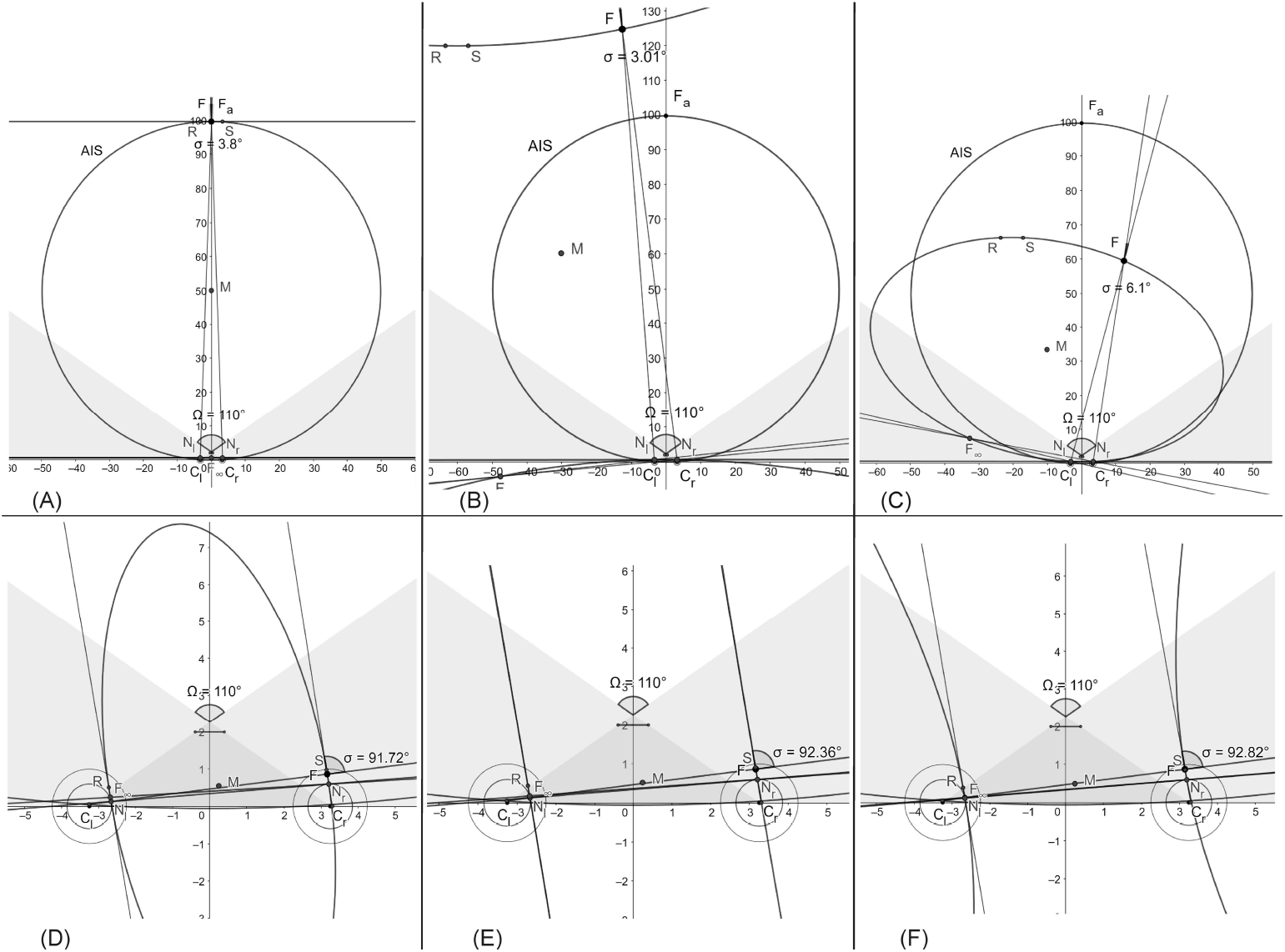
Typical snapshots of the simulation (Supplement 2) in binocular regions are shown in panels (A), (B) and (C). Snapshots in panels (D), (E) and (F) show the atypical conics in the monocular region morphing through two parallel lines from ellipses to hyperbolas at points humans are anatomically unable to fixate on.

To summarize the conclusions of this section, both the perceived direction of the point being fixated on, i.e. the Cyclopean direction, and the binocular conic’s orientation can be assumed to be entities specified by the version angle. This would mean that when the eyes rotate to change gaze in the horizontal plane, both the Cyclopean direction and the binocular conic undergo the same rotations by the version angle. But this would also imply that an object seen to be moving in a frontal line is really moving along the constantly changing horopters in the direction tangential to the instantaneous horopter curves. The curve traced out by the fixation point tracking this object is an iso-subtense curve. This implies that the rotations of both eyes during this pursuit are equal for the AIS passing through the resting vergence position, or that the rotations differ by a constant value along other iso-subtense curves in the horizontal plane. For eyes pursuing other object’s trajectories, the difference in the two eyes’ rotations is time-dependent. For example, this is the case when the object in pursuit moves along the straight frontal line on a flat projection screen in what is a typical laboratory setting.

## 8 Discussion

The horopter’s significance in stereoscopic vision can be explained as follows. When a point in the visual plane lies in front of or behind the horopter curve containing the fixation point, the difference in the angles subtended on each retina between the image and the fovea’s center defines retinal disparity. For each point on the horopter, there is maximum disparity for the single vision that defines Panum’s fusional area around the horopter curve. In this region, noncorresponding retinal elements are fused to provide us with both single vision and the ability to see visual objects stereoscopically in depth from the observer’s current point of fixation. Taking the difference in retinal disparities for a pair of points then provides us with the relative disparity used for our perception of 3D form. Objects outside Panum’s area fall on widely disparate retinal areas and are seen as coming from two different visual directions, causing physiologic diplopia, or double vision. Here, with bicentric projective geometry and a novel model eye, I studied the basic concepts most useful to understanding stereopsic vision: retinal correspondence, horopters and the Cyclopean axis.

### 8.1 Retinal Correspondence and Geometric Horopters

The geometry of longitudinal horopteric conics integrated with eye movements is constructed in the framework of bicentric perspective projections on the image planes of the AEs. The AE is a model eye that includes the eyeball’s global asymmetry caused by the fovea’s displacement from its posterior pole—the main source of the eye’s optical aberrations—and the crystalline lens’ tilt which is countering some of these aberrations [12]. The theory asserts that /*i*/ the longitudinal horopteric curves for the binocular system with AEs are conic sections and /*ii*/, the retinal correspondence obtained from the horopteric conics, is a well-defined concept. Moreover, using this theory allows us to demonstrate that the conics sections’ branches, which pass through the fixation points and referred to as binocular conics, closely resemble empirical horopters obtained by laboratory measurements with the Nonius method [3, 4]. Until recently, there has been only one comprehensive model of empirical horopters and it was elaborated in [5, 6] by an ad hoc introduced equation with a free parameter determined experimentally for each subject. The geometric theory developed here advances that classic model of Ogle and Amigo by establishing a physiologically motivated model of the empirical horopters integrated with the eyes’ movements.

This theory accounts for the fundamental fact that the human visual system functions in physical space and acquires visual information by actively scanning the environment when we are awake. Incident light rays reflected from objects in a scene in 3D space are projected onto the unstable 2D retinae and neuronal processes activated in the visual and visuomotor cortical areas decode and interpret the scene’s 3D properties. Therefore, any decoding of the environment’s 3D properties from sensory information must be fundamentally constrained by the sensory organs’ geometric relationship to the environment [19, 20] and modulated by the eyes’ movements [38].

The conceptual framework used here in constructing horopteric conics for the binocular system with AEs not only provides a biologically-based model that reproduces empirical horopters, it also provides a framework for the theory of geometric horopters developed in [27] for the binocular system with the symmetric (reduced) model eye. This result is proved here in Proposition 1 for any position of the nodal point between the eye’s pupil and its center of rotation, including, of course, the the location of the anatomical nodal point. Thus, three qualitatively different theories of the geometric horopters, including the theory of the Vieth-Müller circle (VMC), are constructed here in the framework of bicentric projections. The three theories are briefly compared below in the order of their respective model eye’s anatomical fidelity.

The first model is a special case of the symmetric model eye in which the nodal point is taken to coincide with the eye’s rotation center. Proposed almost two centuries ago, the resulting horopter curves are the iso-vergence circles, or VMCs, each passing through the fixation point and connecting the eyes’ rotation centers. When the eyes fixate on points along the VMC, the eyes’ rotation centers do not move. This means that the VMC and the vergence value also do not change when the eyes fixate on points along the VMC. Further, relative disparity becomes independent of eye position in this model eye [27]. This model-dependent constancy is a consequence of incorrectly locating the nodal point at a position that is not its anatomical position.

The second model is the symmetric model eye with a nodal point located 0.6 cm anterior to the eyeball’s rotation center as required by the eye’s anatomy. Its horopter curves consist of a family of circles passing through the fixation point and connecting the nodal points [27]. For a constant vergence value, these horopteric circles are parameterized by specific fixation points on the binocularly visible part of the VMC and intersect at the VMC’s point of symmetric convergence. Relative disparity, in this model, depends on eye movement and its changes are always within the binocular acuity limits for fixational eye movements [39]. Regardless of this result, relative disparity is often assumed independent of the eyes’ position. I hypothesized in [27] that the size and shape changes perceived during fixational eye movements may not only provide perceptual benefits, such as breaking camouflage, but may also provide the aesthetic benefit of stereopsis [40].

The third binocular system, with AEs of the highest anatomical fidelity, is the subject of this paper. In this system, the geometric horopters are binocular conics resembling empirical horopters and their orientation is exactly specified by the version angle, giving this angle a new significant meaning in biological vision. On the other hand, if the Cyclopean axis is defined from the midpoint on the VMC’s arc connecting the eyes’ centers of rotation, the same way as it was defined in the binocular system with symmetric eyes, its direction given by the azimuthal angle provides the best approximation of the vergence angle in the human’s binocular system; the difference between the Cyclopean eye direction and the binocular conic orientation given by the version is on the order of a few seconds of an arc (Figure 5).

Although the VMCs and empirical horopters have different geometries, the VMC is often identified with the longitudinal horopter. The VMC does provide a good approximation for the empirical horopter near the fixation point, but the difference in their geometries is significant in the periphery. A small object peripherally located on the VMC will have zero disparity with respect to this horopter model, but it will have a nonzero disparity with respect to the binocular conics that approximate well the empirical horopters over the whole visual field. Visually guided saccades intercepting a peripherally viewed object will be well off the target if programmed in terms of VMC’s disparity. Although the simplicity of the VMC makes it useful in some numerical aspects of visuomotor research, its approximation of both geometric and empirical horopters is a crucial condition that should always be emphasized in order to avoid its, currently frequent, mischaracterizations.

Further, it is suggested in [41] that the shape of the longitudinal horopter is a result of the visual system allocating resources according to natural disparity statistics for binocular correspondence matches. Although the horopter’s shape can support these statistics, my theory instead asserts that the shape of empirical horopters is caused primarily by the misaligned optical elements modeled by the AE. In fact, in healthy eyes, the fovea is displaced from the eyeball’s posterior pole and the cornea and the crystalline lens are tilted relative to each other [9, 10, 23]. The crystalline lens’ tilt cancels out some of the aberrations caused by foveal displacement and the cornea’s asphericity and produces nearly aberration-free perception near the visual axis [11, 12]. Then, the adaptation to the natural environment’s visual statistics can be achieved through the binocular eye’s movements [35, 42].

### 8.2 Binocular Conics and Eye Movement

The fovea, which has the highest visual acuity on the retina, subtends only a two-degree visual angle. To prevent diplopia, a saccade must quickly direct the eyes’ foveae toward the object—in what is called the conjugate eye movements because the eyes are rotating in the same direction. Saccades usually need to be corrected by a vergence—the disjunctive eye movements as they rotate in opposite directions, and then the foveae must be held precisely aligned on the object [43, 44]. This corrective vergence movement, or motor fusion, adjusts the eyes’ alignment to maintain sensory fusion [45, 46].

Cortical activity derived from bicentric perspective retinal stimulation must, therefore, be modulated by the eyes’ movements [38]. The size and direction of the adjustment is given by the binocular disparity between the currently viewed object and the next one to be viewed. Thus, the concept of retinal corresponding elements is not only fundamental to single vision and stereopsis, it is also important in the binocular coordination of the eyes’ movements. Understanding how the eyes’ movements are controlled by the visuomotor processes and how they affect the precise correspondence of the retinal elements remains uncertain [47].

Moreover, during natural viewing, the human eye’s rotational speeds during saccades are as fast as 700°*/*s, with an acceleration exceeding 20, 000°*/s*^2^ [47]. Saccadic eye movements are performed about 3-4 times/s, meaning that visual information is mainly acquired by the brain during 3-4 brief fixations within a second. In addition, we are not only able to execute smooth pursuit eye movements that keep the foveae focused on a slowly moving object up to 100°*/*s, we also employ a combination of smooth pursuit and saccades to track an object moving unpredictably or moving faster than 30°*/*s [48, 49]. By stabilizing the tracked object’s image on the fovea, smooth pursuit eye movements (SPEMs) superimpose additional motion on the retinal images of the stationary background and on the moving objects.

For example, the consequences of the saccadic eye movements’ high speed and acceleration markedly restricts the use of visual information between fixations. Therefore, the basic feature underlying natural viewing is the occurrence of intricate dynamic disparity that is then processed to maintain our clear vision that appears continuous and stable. In this regard, my theory provides the binocular conics’ transformations by integrating the binocular conic’s geometry with the eyes’ changing position in the horizontal visual plane of bifoveal fixations, therefore extending my work on modeling the monocular vision stability in [50, 51] to the binocular framework.

The kinematics of visually guided eye movements is constrained by Listing’s law which involves the primary eye position in this law’s formulation. In its typical version, which originally applied to a single eye’s rotation, when the eye fixates on a target at optical infinity, Listing’s law asserts that, with the head upright and stationary, there is an eye position called the primary position such that any other eye orientation can be reached by a single eye rotation about the axis in the plane perpendicular to the eye’s primary direction. This plane is known as Listing’s plane. Consequently, during eye movements that obey Listing’s law (e.g., saccades and smooth pursuit), the eyeball assumes a unique torsion, or a rotation about the line of regard, for each eye orientation [52]. In my study, all eye movements are constrained to rotate about the vertical axis such that the torsion specified by Listing’s law is always zero.

Further, when both eyes are constrained to fixate binocularly during the eyes’ rotations, binocular extension of Listing’s law, known as L2, applies to the eyes’ positions. L2 asserts that during convergence, the eyes’ rotation axes still remains confined to a plane for each vergence angle; however, as the eyes converge, these planes rotate relative to Listing’s planes temporally and roughly symmetrically [53, 54, 55].

My theory specifies the AEs’ primary position at the abathic distance fixation, or the resting vergence position, to allow for the physiologically motivated replacement of the imprecise primary eye position as follows. In the absence of visual cues, the eyes’ gaze shifts to the eyes’ natural tonus resting vergence position, which serves as a zero-reference level for convergence effort [17]. In fact, the average natural tonus resting vergence distance for the forward gaze is of the same value as the human’s average abathic distance, which also agrees with the abathic distance for the average anthropomorphic parameters of the AE. Moreover, though both the tonus resting vergence distance and the abathic distance vary from about 40 cm to optical infinity across subjects, they are reliable parameters within a subject [56]. Further, the resting vergence position is supported by recent results demonstrating that Listing’s law and kinematics related to Listing’s law are implemented peripherally and by the oblique extraocular muscles (EOMs) mechanism rather than centrally [57]. In fact, these EOM’s forces are indispensable to 3D modeling of eye movements and are responsible for the mechanical equilibrium of the eye suspended in resting vergence position [58]. Moreover, the resting vergence position’s change with lowered and elevated gaze [59] agrees with the vertical horopter’s backward inclination and its effect on perception [60, 61, 34].

The above discussion strongly support my choice of the resting vergence position at the abathic distance of bifoveal fixation to replaces the eyes’ primary position that essentially applies only to a single eye, but is often stated as describing both eyes fixating at optical infinity with an obvious lack of precision. This could be the reason that despite theoretical importance of the eyes’ primary position, its precise formulation and neurophysiological significance remain elusive [18].

The theory of binocular conics constructed here needs to be further extended with the vertical component and integrated with 3D eye movements. This extension to a full framework of bicentric perspective [16] will inevitably introduce a host of geometric difficulties. For example, the visual line in the AE model that passes through the fovea, the optical node, and the fixation point differs, because of the fovea’s displacement from the posterior pole of the eyeball, from the line of regard, or fixation axis, connecting the eye’s rotation center with the fixation point. One of the questions this introduces is how to model the rotation of the visual axis by the eye’s torsion around the line of regard because a rotation complicates the control of the binocular eyes’ alignments in near-vision conditions. Moreover, the extension of Listing’s law which applies when the eyes start rotating from their tertiary position, the so called half-angle rule, was not needed here, but will be indispensable when my theory is extended to 3D rotations because this extensions requires the use of angular velocity rather than rotation axes [62].

Also specific to 3D kinematics of eye-head bifoveal fixations is either the Listing’s plane’s tilt or Listing’s plane geometry change to a twisted surface. The Listing’s plane’s geometry changes have been analytically modeled as the effect of the alignment maximization method [63]. Further, it has been proposed in [64, 65] that the crystalline lens’s horizontal tilt and the eye’s axial length change during a 25° downward gaze with binocular fixations at the 0.2 D and 2.5 D accommodative states for near-visual tasks. Although recent results in [66, 67] support this proposition, the theory regarding the physiological mechanism of accommodation is still incomplete. It appears that the crystalline lens can change its tilt due to a loosening of the zonule—the fiber band attached to the lens that changes its curvature during accommodation—and gravity [68], and careful modeling of this and other accommodative mechanisms may contribute to a fuller understanding of the origins of the elusive presbyopic changes [69]. Thus, modeling physiological mechanisms underlying stereoscopic vision should include not only the eye’s optical asymmetry but also the tilted, accommodating crystalline lens.

## Acknowledgements

I thank Alice Turski for her editing that has made this manuscript much easier to read.

## Supplemental Materials

### Supplement 1

The proof of Proposition 1 in Section 5 uses results obtained in [1], which are shown in Figure S1 below. The large circle drawn in a dash-dot line is the Vieth-Müller circle (VMC) with its center *C*_*V*_ and the large circle drawn with a solid line is the geometric horopter circle (GHC) with center *C*_*H*_. The vergence is *η* = *ϕ*_*r*_ *− ϕ*_*l*_ and the version is *ω* = 1*/*2(*ϕ*_*r*_ + *ϕ*_*l*_). The point *S* of symmetric convergence is the intersection of the VMC with the GHC.

**Figure S1.**
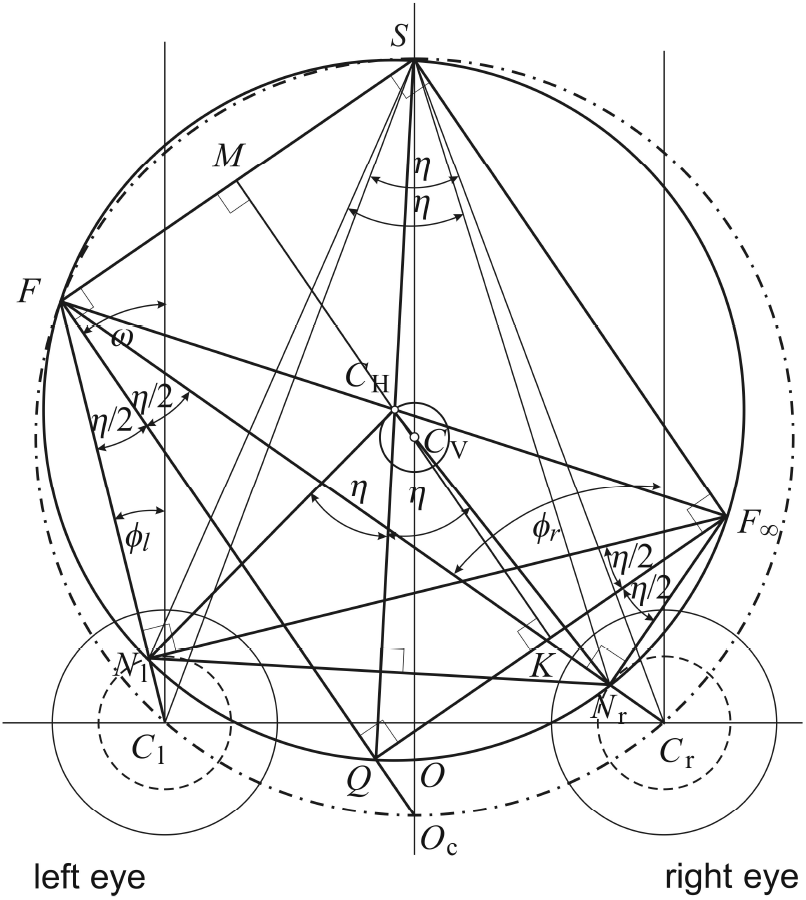
Proof of Proposition 1

#### Proposition 1 1.

*Let the nodal point be located on the optical axis at any point at or between the spherical eyeball’s rotation center and its pupil. Then, for the binocular eyes position with fixation point F in the horizontal visual plane, the lines passing through the nodal points and perpendicular to the visual axes intersect at the point F*_*∞*_ *on the circular horopter. It then follows that line segment FF*_*∞*_ *must pass through the horopteric circle’s center*.

*Proof*. When the eyes are fixated on *F*, the GHC is defined by three points: *F* and the two nodal points, *N*_*r*_ and *N*_*l*_. According to the Central Angle Theorem, the rays perpendicular to the visual axes at the nodal points intersect at point *F*_*∞*_ on the GHC. This is demonstrated by the triangles *QN*_*l*_*F* and *QN*_*l*_*F*_*∞*_ in Figure S1 sharing the same angle at vertices *F* and*F*_*∞*_. Then, by results proved in [1], line segment *FF*_*∞*_’s midpoint *C*_*H*_ is the center of inscribed rectangle □*QF*_*∞*_*SF* and, therefore, also the center of GHC’s center. This completes the proof.□

### Supplement 2

#### A. Binocular Conics Transformation

The *GeoGebra* applet BCT1 at https://www.geogebra.org/m/m8btzc5w allows us to visualize the binocular conics’ (shown in red) transformations in the visual plane of bifoveal fixations. The initial abathic-distance fixation *F* has coordinates (0, 99.61). The initial conic at this resting vergence position consists of two parallel horizontal lines where the line passing through *F* is the linear horopter. When point *F* moves, the window ResetF displays the point *F* ‘s current coordinates. After the session is finished, return *F* to the position on the intersection of the *y*-axis and the AIS circle that locks to (0, 100), so reset coordinates in ResetF from (0, 100) to (0, 99.61) to retur*F* to the resting vergence position.

1. Open the above link in your browser.
2. In the applet, click on the red *F* of the resting vergence position at (0, 99.61) to highligtht the red dot.
3. Drag the red point with your mouse through the visual plane to transform the conics.
4. Return *F* to the resting vergence position (described above) by resetting coordinates in ResetF to (0, 99.61).

#### B. Retinal Correspondence

Remark 1 in Section 5, asserts that the retinal correspondence defined in **Retinal Correspondence** in Section 3 is well defined concept. I demonstrate here this assertion with *GeoGebra*’s simulation of binocular conics transformation. In the*GeoGebra* applet BCT, the point *Q* on the linear horopter (shown in blue) is projected along visual lines to points *Q*_*r*_ and *Q*_*l*_ on the right and left image planes, respectively. The angles between these visual lines of point *Q* and the corresponding initial visual axes of *F* are displayed as *ϕ*_*Qr*_ and *ϕ*_*Ql*_ for the right and left eye respectively. When the resting vergence position fixation point *F* is moved (cf. the previous subsection), the value of these angles unchanged to 3 decimal places. This proves that the retinal correspondence relationship defined in Section 3 of the article is invariant of eyes position in the visual plane of bifoveal fixations and, therefore, is a well-defined concept.

